# A generalization of the informational view of non-random mating: Models with variable population frequencies

**DOI:** 10.1101/279075

**Authors:** A. Carvajal-Rodríguez

## Abstract

Mate choice may generate non-random mating patterns. It has been recently shown that the mating distribution caused by mate choice can be expressed as a gain in information with respect to random mating. In that model, the population phenotypic frequencies were assumed as constant during the breeding season. In the present work such restriction was relaxed to consider different encounter-mating processes in which the population frequencies of available individuals change over mating rounds. As with the constant case, here we describe the change in the mating phenotypes by the flow of information with respect to random mating. This information can be partitioned into sexual selection, sexual isolation and a mixed effect. Likewise, the pairwise statistics for total change, sexual selection and sexual isolation are generalized for variable population frequencies.

The new tests had more power for the detection of the effects of non-random mating when the population frequencies vary during the breeding season. The differences in power were high for sexual selection but slight for sexual isolation scenarios. However, the application of the new formulas require the estimation of frequencies at each mating round. Therefore, choosing one or another type of statistics would depend on the biological scenario as well as in the availability and easiness to split the sampling in more than one mating round.

## 1. Introduction

Mate choice can be defined in general, as any aspect of an organism phenotype that leads to an individual to engage in sexual activity with some partners more likely than with others (Rosenthal, 2017).

From a population genetics point of view, mate choice is defined by its effects: it is the observed mating frequency deviation with respect to random mating. So defined, the effects of mate choice can be partitioned into sexual selection and sexual or behavioural isolation (intersexual selection). Sexual selection is defined as the observed change in gene or phenotype frequencies in mated individuals with respect to population frequencies. Sexual isolation is defined as the deviation from random mating in mated individuals (Rolán-Alvarez and Caballero, 2000).

In a previous work (Carvajal-Rodríguez, 2018), the mating distribution caused by mate choice was expressed as the gain in information with respect to random mating. In that model, the population phenotypic frequencies were assumed as constant over the breeding season. In the present work, the previous results were extended to include the case where population frequencies vary over different mating rounds during the breeding season.

The extension to include variable population frequencies is appropriate for a better description of monogamous species in which pair formation occurs by different mating rounds. In this case, the frequencies of available individuals can change during the same breeding season (Gimelfarb, 1988).

We show that, by relaxing the condition of constant population frequencies, a new generalized information partition can be obtained. As before, this information can be partitioned into sexual selection, sexual isolation, and a mixed effect, while the corresponding G-like tests can be performed over each information index (Carvajal-Rodríguez, 2018). The new equations become the previous ones as soon as the population frequencies are set to constant values.

Prairie vole (*Microtus ochrogaster*) is a good model organism that can help us to demonstrate the new tests. This mammal is a monogamous species that establishes long-term heterosexual relationships among adult mates. This rodent lives mainly in the grasslands of the central United States and Canada. Its living conditions imply limited food and water resources, which may have contributed to the evolution of a monogamous life strategy in this species (Williams et al., 1992; Young et al., 2011). It is known that the long-term pair-bonds in this species are regulated by a variety of neurotransmitters and driven by epigenetic events (Wang et al., 2013). Indeed, the prairie vole has become a model system for the study of the neurobiology of monogamy, social attachment and nurturing. However, for our purpose is enough to consider that, after encounter and mating, there is an oxytocin-modulated monogamous behaviour which, from the point of view of the frequency of available adults, may resemble a model without replacement. That is, the already formed mating pairs are excluded from the mating pool. Therefore, we simulated sampling in each of several mating rounds to compare variable versus constant frequency tests for three different scenarios (random mating, sexual isolation and sexual selection).

Following, we first obtained the general information equation for the total change, and then developed the partition into the different effects. Afterwards, the new model was demonstrated by example, and finally the new outcomes were summarized in the discussion.

## 2. Non-random mating with variable population frequencies

During the breeding season the formation of mating pairs is a two-step process in which first, an encounter occurs and second, it ends or not in a successful mating (Gimelfarb, 1988).

When males and females can have multiple mates, or when only a fraction of the individuals of both sexes do actually mate in the current breeding season, it is reasonable to assume that the availability of the different phenotypes are not affected by the matings already occurred. Therefore, the population frequencies can be considered constant during the reproductive season. In this case, the process of mating resembles a sampling with replacement from the point of view of the available phenotypes (Carvajal-Rodríguez, 2018).

On the other hand, when we consider a monogamous species in which most of the sexually mature individuals are involved in a series of mating rounds, then the availability of individuals is affected by the previous matings. In this case, the population phenotypic frequencies are not constant during the breeding season and the process resembles a sampling without replacement.

### 2.1 General model

Let consider a population where the mating occurs in various sequential rounds so that in the *r*^*th*^ round the number of available (unmated) females is *n*_1r_ and the number of available males is *n*_2r_. Thus, the relative frequency of the female phenotype *i* is *p*_1ri_ = *n*_1ri_ / *n*_1r_, i.e. the number of available *i* - type females divided by the total number of available females. If the model is without replacement, the frequencies may be different in the different mating rounds. Similarly, for males with phenotype *j* we have *p*_2rj_ = *n*_2rj_ / *n*_2r_.

During the breeding season, the absolute frequencies of adults will be updated depending on the type of the encounter-mating scheme. Under the individual encounter-mating model (Gimelfarb, 1988) there is only one mate by round. In this case, if the new mating includes a female of *i* - type then in the next round we have *n*_1(r + 1)i_ = *n*_1ri_ - 1 and *n*_1r + 1_ = *n*_1r_ - 1, and similarly for males. In general, if each mating round involves a number *U*_*r*_ of matings then in the next round, we will have *n*_1r + 1_ = *n*_1r_ - *U*_*r*_ available females and *n*_2r + 1_ = *n*_2r_ - *U*_*r*_ available males, and the corresponding phenotypic classes updated depending on the specific matings that occurred.

Now, consider the encounter of a female of type *i* with a male of type *j*. The mutual mating propensity *m*_*ij*_ is the number of matings, after encounter of *i* with *j*, in a given environment (Carvajal-Rodríguez, 2018).

In mating round *r* there is a probability *q*_rij_ = *p*_1ri_ × *p*_2rj_ of encounter between *i* and *j*. Therefore, the number of matings *i* × *j* at this round is *q*_rij_*m*_ij_. The sum over the mating rounds gives the total number of *i* × *j* matings and so, the observed relative frequencies of these matings after *R* rounds can be expressed as

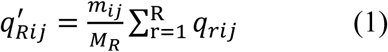

where 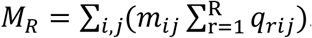.

Note that *M*_*R*_ defines the total number of matings at the end of the season and can be expressed as the sum of matings at each round so that

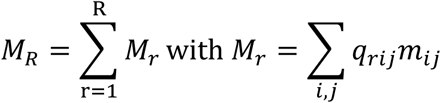

Also note that the maximum possible number of matings occurring in round *r* is *max*(*M*_*r*_) = *max*{*m*_*ij*_, …} for every *i* and *j*.

When mating is at random, the mating propensity is constant i.e. *m*_ij_ = *m* for every pair *i, j* and so

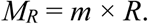

Then, by noting the expected frequencies under random mating as *q*_*R*îj_ and by substituting the constant propensity in (1) we obtain

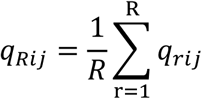

Note that if the frequencies are constant over the different mating rounds then we recover the formulas as given in (Carvajal-Rodríguez, 2018). So that with constant frequencies

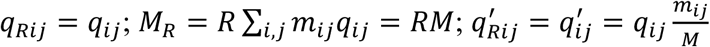

When the population frequencies are variable over the breeding season, the correct pairwise statistic for measuring the deviation from random mating is given by RPTI_ij_ = *q*’_*R*ij_/*q*_*R*ij_, which is the ratio of the frequency of the observed pair types divided by the expected pair types under random mating, calculated from the population frequencies when these are variable over the same breeding season. If the population frequencies can be considered constant, the RPTI_ij_ statistic becomes the pair total index (PTI_ij_) as originally defined in (Rolán-Alvarez and Caballero, 2000).

As with the constant frequency case we can measure the change with respect to random mating

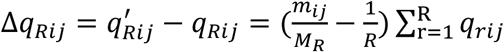

and from (1) we can also express the propensity as a ratio of frequencies

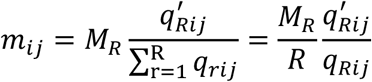

to finally obtain the expression for the mean population change in terms of the logarithm of the mating propensity as

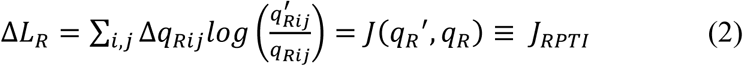

which is the Kullback-Leibler symmetrized divergence that measures the gain in information when the differential mating propensity moves the population from mating frequencies *q*_*R*_ to *q*’_*R*_ or vice versa. Note that *J*_*RPTI*_ becomes the information equation for the constant case when the population frequencies are set to constant values (*J*_*PTI*_ in eq. (2) in Carvajal-Rodríguez, 2018).

As with *J*_*PTI*_, if we take the natural logarithm in (2) and multiply by the total number of matings, the obtained quantity is well approximated by a chi-square under the null hypothesis of random mating. The possible difficulty for performing this test, is being able to estimate the expected frequencies under random mating over the distinct mating rounds.

### 2.2. The information partition

To perform the information partition, we have to take into account that the expected pair types under random mating calculated from mated individuals, must be computed within each mating round.

In round *r*, the number of matings involving females of phenotype *i* is

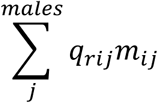

Then, the relative frequency of females *i* calculated from the matings within this round is

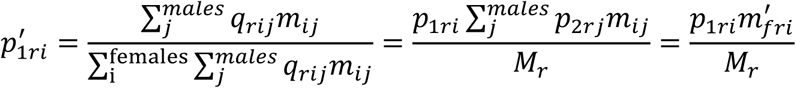

where 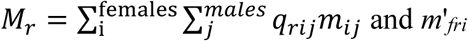 is the marginal propensity of females with phenotype *i* in the round *r*.

Similarly, the frequency of males calculated from the matings,

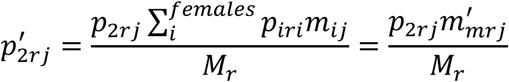

where *m*’_*mrj*_ is the marginal propensity of males with phenotype *j* in the round *r*. Also, note that Σ_i_*p*’_1ri_ = *M*_*r*_ / *M*_*r*_ = 1 and Σ_i_*p*’_2r_ = *M*_*r*_ / *M*_*r*_ = 1.

Then, after *R* rounds we get the probability of random mating *i* × *j* computed from matings as

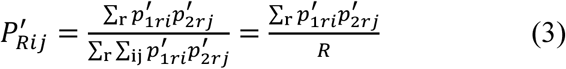

Now, we may express the components of *J*_*RPTI*_ in (2) as

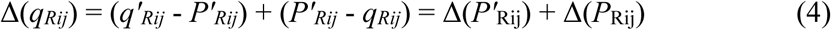

and

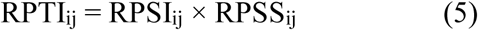

where RPSI_*ij*_ = *q’*_*Rij*_ / *P’*_*Rij*_ and RPSS_*ij*_ = *P’*_*Rij*_ / *q*_*Rij*_ correspond to the generalized version of the pair sexual isolation and pair sexual selection statistics respectively, for the case of variable population frequencies over different mating rounds. These statistics become the original ones when the population frequencies are constant.

By substitution of (4) and (5) in (2) we obtain the information partition

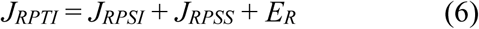

where

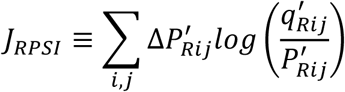

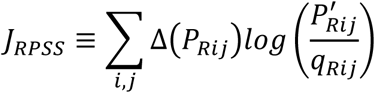

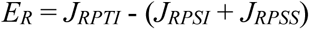

We have generalized the information partition of total change due to non-random mating, for models without replacement in which the population frequencies are updated after each mating round. The original partition developed in (Carvajal-Rodríguez, 2018), emerges as a particular case from (6) when the population frequencies are constant or the number of mating rounds are reduced to one (*R* = 1). Concerning the sexual selection information, in the previous work we showed that under the constant frequency assumption, sexual selection can be divided into its female and male components. However, the generalized index (*J*_*RPSS*_), is no longer the exact sum of the information in both sexes. There is a new error term that increases with the number of mating rounds. To appreciate this, let consider the female and male population frequencies averaged over *R* mating rounds

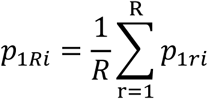

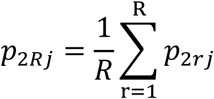

And the averages for the frequencies calculated from mated individuals, i.e.

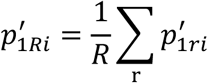

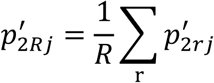

Now we define the sexual selection effect in females

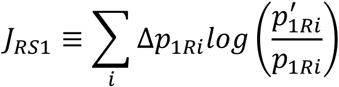

and in males

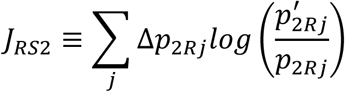

where Δ*p*_*1Ri*_ = *p’*_1Ri_ - *p*_*1*Ri_ and Δ*p*_*2Rj*_ = *p’*_*2Rj*_ – *p*_*2Rj*_.

Then, the sexual selection information is

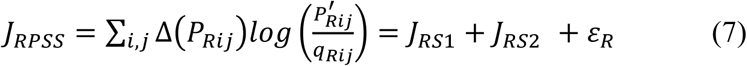

the error term arises because the logarithm for *P’*_*Rij*_ and *q*_*Rij*_ correspond to a sum of products over *R*, which makes sense because encounters only happens within the same round i.e., for *r* ≠ *s*, the quantities *p*_1*ri*_ × *p*_2*sj*_ or *p*_1*si*_ × *p*_2*rj*_ have no biological sense here. For the partition in (7) to be exact (*ε*_*R*_ = 0) we need to equate the sum of products to the product of the sums, i.e. Σ_*r*_ (*p*_1*ri*_ × *p*_2*rj*_) = (Σ _*r*_ *p*_1*ri*_) × (Σ _*r*_ *p*_2*rj*_) which is not true in general. The same problem occurs with *p’* _1*ri*_ and *p’* _2*rj*_. The discrepancy increases with the number of rounds while it disappears if we have only one round or the frequencies are constant.

There is still another issue with the information partition estimates under the individual encounter models. The equation (3) corresponds to the formal model for the expected random mating calculated from matings instead of from population frequencies. It is usually estimated simply by the relative frequency of phenotypes within the matings at each round. However, under the individual encounter model there is only one mating per round, and estimating (3) by the count of one individual in the mating per round, produces a value of *P’*_*Rij*_ that is exactly the same as *q’*_*Rij*_ from equation (1) so that, the sexual isolation effect is not detected but included as a sexual selection effect. In fact, encounter models where the number of mating per round is equal or less than the product of the number of phenotypic classes (*k*_1_ × *k*_2_), will be biased against the detection of sexual isolation effects if the estimation of *J*_*RPSI*_ is performed by counting matings per round. A solution for this problem will be suggested in the Example section below. On the contrary, when the number of matings per round is higher than the product of phenotypic classes (e.g. *k*_11_ × *k*_2_ = 4) the estimation works better than the constant frequency estimators.

### 2.3. Generalized joint isolation index

Having defined RPSI_*ij*_ as a generalized version of the sexual isolation pairwise statistic for variable population frequencies, we can obtain the corresponding joint isolation index as

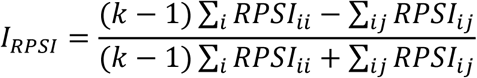

where *k* is the number of phenotypic classes (*k* = *k*_1_ = *k*_2_ for the joint index to be applicable). The index matches the original one as defined in (Carvajal-Rodriguez and Rolan-Alvarez, 2006) when the population frequencies are constant.

When the population frequencies are constant (models with replacement), we already know that intersexual selection (sexual isolation) emerges as a deviation of the multiplicative effect in the propensities. That is, if the normalized mutual propensity is the product of the relative marginal effects of each partner there is no sexual isolation (Carvajal-Rodríguez, 2018). However, in the general model the condition is affected by the mating round and it becomes

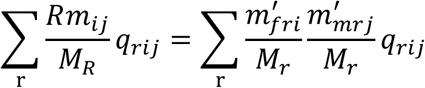

so that, a sufficient condition for the absence of sexual isolation effects over the frequencies is

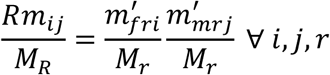

Note that under constant frequencies *m*’_*fri*_ = *m*’_*fi*_; *m*’_*mrj*_ = *m*’_*mj*_; *M*_*r*_ = *M* and *M*_*R*_ = *R* × *M* and then we recover the same condition as from formula (5) in (Carvajal-Rodríguez, 2018).

## 3. Example

The difference between the variable frequencies and the constant frequencies approaches can be shown by a toy-model based on a monogamous species. One of these species is the prairie vole (*Microtus ochrogaster*) that is a monogamous rodent that establishes long-term relationships between adult mates.

Concerning mate choice, there is no clear evidence of prairie vole mate choice behaviour in field studies. Indeed, some results indicate that prairie vole just mate with the first available partner (Getz et al., 2004; Keane et al., 2007). On the other hand, laboratory studies demonstrate that length polymorphism of microsatellite DNA related with genes within the vasopressin–oxytocin pathway (e.g. arginine vasopressin receptor 1a, *Avpr1a*, and oxytocin receptor, *Oxtr*) are correlated with female social and sexual preferences (Castelli et al., 2011; Keane et al., 2017). In addition, a potential for assortative mating at the *Oxtr* locus has recently been suggested in the bank vole (*Myodes glareolus*) species (Watts et al., 2017).

In consequence, we simulated three examples, first for a trait not linked with choice (random mating), and second and third, for a trait with positive assortative or sexual selection effects. For each example, we assayed both an individual and a mass encounter models (Gimelfarb, 1988). In the first, only one pair mate at each mating round while in the second up to 10 individuals can mate simultaneously. In both cases, the population frequencies were updated after each mating round.

The population consisted of 200 sexually mature adults with 50 individuals of each phenotype and sex. The species is monogamous, and assuming the whole population is involved in the breeding season, we can have up to 100 mating pairs. Therefore, we need 100 mating rounds (*R* = 100) to sample 100 matings under the individual model (just one mating per round) while we need at least 10 rounds (*R* ≥ 10) for sampling the same number of matings under the mass encounter model (at least one mating and a maximum of 10 at each round).

For comparing the performance between the two types of information indices we ran 10,000 breeding seasons for each case.

### 3.1. Random mating

As a trait not related to choice, we considered the belly which in this species is yellow, ranging from pale to dark, and it has not known effect on mate choice. Thus, we may classify the matings by the belly color phenotype as pale yellow (*y*) or dark (*d*). The initial frequency of the phenotype was assumed to be equally distributed with intermediate frequency in both sexes so *p*_1*y*_ = *p*_2*y*_ = 0.5 and *p*_1*d*_ = *p*_2*d*_ = 0.5.

Because mating is at random (mutual propensities are equal) we simply performed the Mont Carlo sampling based on the phenotypic frequencies. That is, to incorporate a female into the mating pair, we checked if a uniform *U*(0,1) random number was lower than *p*_1*y*_ and then a yellow female entered to the pair, if not, a dark female did it. For incorporating the male partner, we checked if a new random number was lower than *p*_2*y*_ then a yellow male also entered to the pair, if not, a dark male did it. After 1 (individual model) or 10 (mass model) pairs were formed, the frequencies were updated, and the process repeated until a total of 100 pairs was reached.

In Table 1, we can see the result of 10 mating rounds under the mass encounter model. During the mating rounds, the population frequencies were recorded jointly with the number of matings for the different classes. At the end of the breeding season (after round 10), the sum of the observed matings over rounds corresponded to the finally observed matings that we would have recorded if we do not distinguish among the different mating rounds.

**TABLE 1.** Result of 10 mating rounds under mass encounter with random mating: *M*_*yy*_ = *m*_*yd*_ = *m*_*dy*_ = *m*_*dd*_ = 10.

| Round $r$ | $p_{1ry}$ | $p_{2ry}$ | $y \times y$ | $y \times d$ | $d \times y$ | $d \times d$ |
| --- | --- | --- | --- | --- | --- | --- |
| 1 | 0.5 | 0.5 | 1 | 4 | 4 | 1 |
| 2 | 0.5 | 0.5 | 3 | 3 | 4 | 0 |
| 3 | 0.487 | 0.475 | 5 | 2 | 1 | 2 |
| 4 | 0.457 | 0.457 | 5 | 2 | 1 | 2 |
| 5 | 0.417 | 0.433 | 0 | 2 | 2 | 6 |
| 6 | 0.46 | 0.48 | 4 | 1 | 3 | 2 |
| 7 | 0.45 | 0.425 | 1 | 0 | 5 | 4 |
| 8 | 0.567 | 0.367 | 1 | 5 | 1 | 3 |
| 9 | 0.55 | 0.45 | 4 | 3 | 0 | 3 |
| 10 | 0.4 | 0.5 | 2 | 2 | 3 | 3 |
| Observed total |  |  | 26 | 24 | 24 | 26 |
| | | $q'_R$ | 0.26 | 0.24 | 0.24 | 0.26 |
| | | $q_R$ | 0.219 | 0.26 | 0.24 | 0.281 |
| | | $\Delta q_R$ | 0.04 | -0.02 | 0 | -0.02 |
| | | $\ln(q'_R / q_R)$ | 0.172 | -0.08 | 0 | -0.078 |
| $J_{RPTI}$ (p-value) | 0.01 (0.79) | | | | | |
| $J_{PTI}$ (p-value) | 0.002 (0.98) | | | | | |

$q'_R$ : observed mating frequencies; $q_R$ : expected random mating frequencies computed following the model in equation (1) by taking equal propensities; $\Delta q_R$ : $q'_R - q_R$ . $\ln$ : natural logarithm.

With the recorded data, we can compute *J*_*RPTI*_ and *J*_*PTI*_ which for the specific case in the Table 1 gave non-significant values at 5% significance level (*J*_*RPTI*_ = 0.01, *J*_*PTI*_ = 0.002). Recall that each test is performed multiplying the corresponding index by the total number of matings. The obtained value follows a chi-square distribution with 3 degrees of freedom under the null hypothesis of random-mating.

This mating process was repeated 10,000 times and the obtained false positive rate was 4% for *J*_RPTI_ and 1% for *J*_PTI_. The same percentages were obtained for the individual encounter model.

### 3.2. Non-random mating: Sexual isolation

As a putative assortative mating trait, we considered the length polymorphism of microsatellite DNA related with *Oxtr*. Thus, we may classify the matings by long (*L*) versus short (*S*) length polymorphism. The population size, mating rounds and initial frequencies were defined as in the previous example, so, *p*_1*L*_ = *p*_2*L*_ = 0.5 and *p*_1*S*_ = *p*_2*S*_ = 0.5. As before, there would be 100 mating rounds under the individual encounter model and at least 10 mating rounds under the mass encounter model. In the latter, there are 10 encounters per round but some of them may not succeed, so more than 10 mating rounds would be necessary for completing 100 matings.

To model assortative mating, the mutual propensities were different and we defined the homotypic matings having three times more propensity than the heterotypic, so *m*_*LL*_ = *m*_*SS*_ = 3*m*_*LS*_ and *m*_*SL*_ = *m*_*LS*_. The specific value of the maximum propensity depended on the encounter model i.e. *m*_*LL*_ = 1 for the individual encounter and *m*_*LL*_ = 10 for the mass encounter model.

### 3.2.1 Mass encounter

In each mating round, 10 random encounters occurred based on the population frequencies. Once the encounter happened, the mating succeeds if a uniform *U*(0,1) random number was lower than the pair propensity normalized by *M*_*r*_, if so, the mating was recorded. The frequencies were updated after the 10 encounters. As in the random mating case, the simulation ended when 100 matings pairs were formed.

With the recorded frequencies we computed *J*_RPTI_ and *J*_PTI_, and performed the corresponding tests, which for the specific mass encounter example shown in Table 2, were significant for *J*_RPTI_ (*J*_RPTI_ = 0.13, *p* - value = 0.004) but non-significant for *J*_PTI_ (*J*_PTI_ = 0.04, *p* - value = 0.256).

**TABLE 2.**
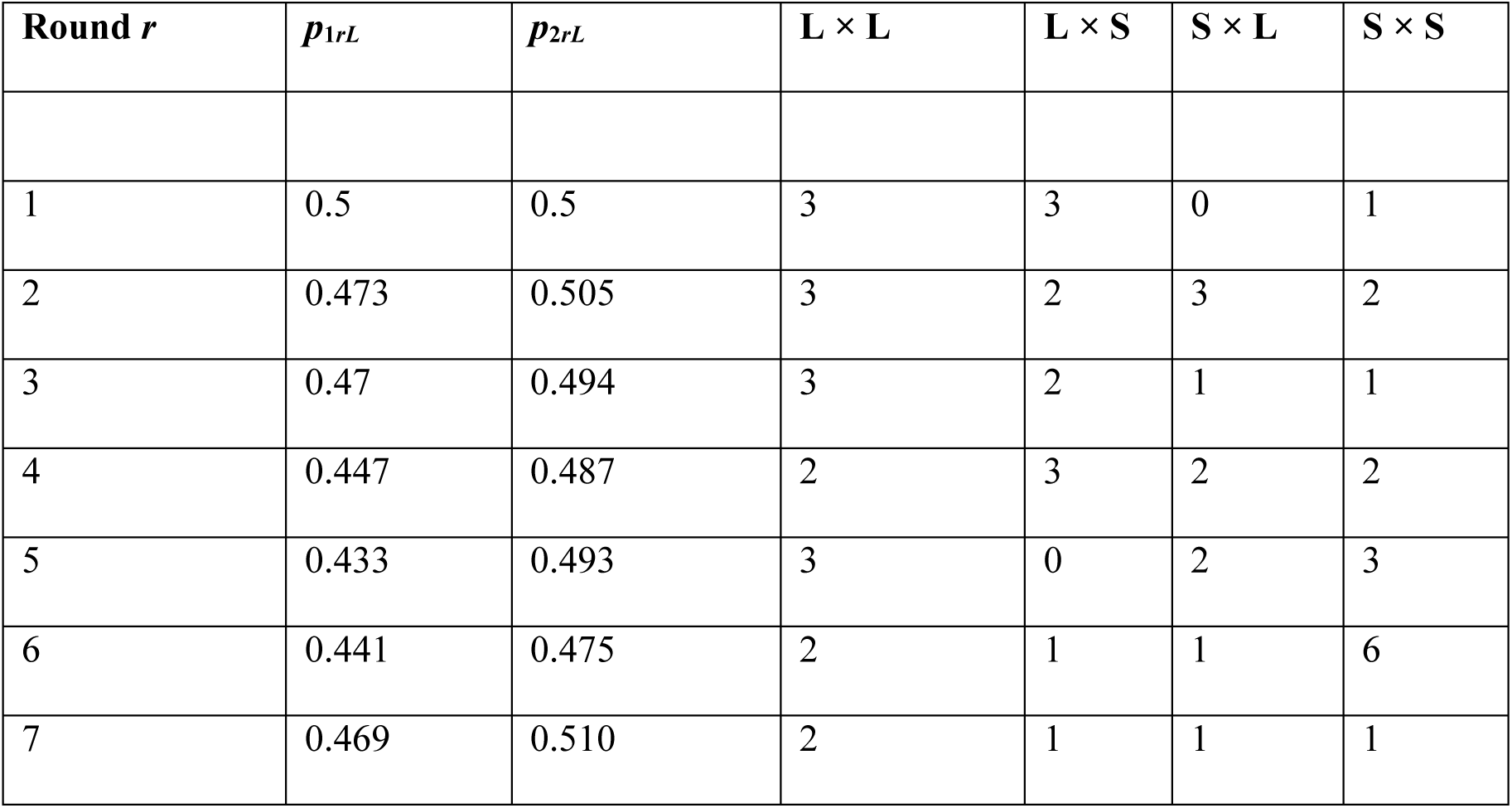
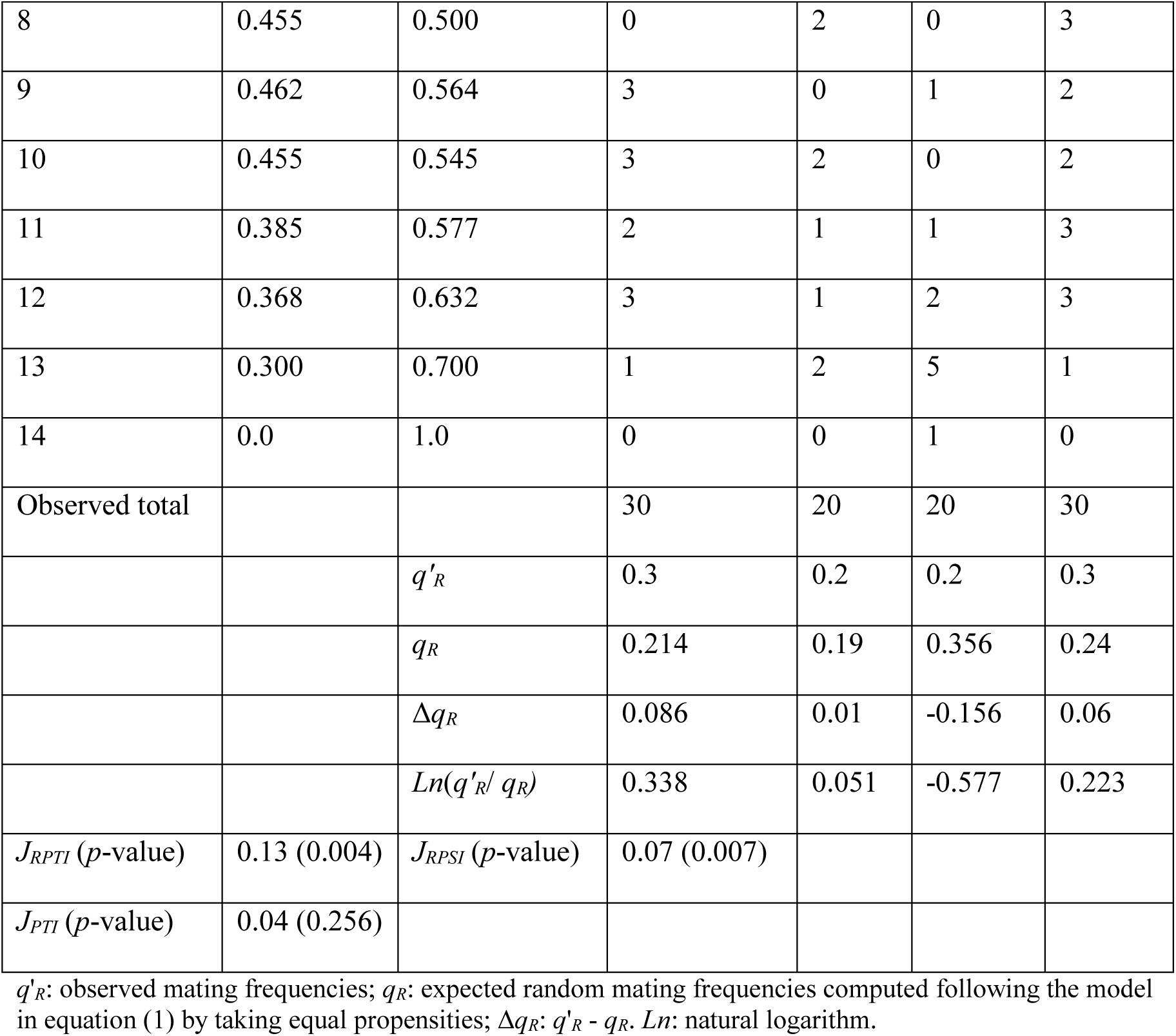
Result of 14 mating rounds under a mass encounter model with assortative mating: *m*_*LL*_ = *m*_*SS*_ = 10 = 3*m*_*LS*_, *m*_*SL*_ = *m*_*LS*_.

In general, the power for detection of non-random mating was about 90% for *J*_RPTI_ and 73% for *J*_PTI_ (over 10,000 runs). In the case of detection of sexual isolation effects, the *J*_RPSI_ test had a power of 87% while a remaining 3% corresponded to the miss-detection of sexual selection (*J*_RPSS_) instead of isolation. However, in the case of *J*_PSI_ all the non-random mating detected cases were identified as sexual isolation (73%).

### 3.2.2 Individual encounter

In the case of the individual encounter, the differences between the indices for detection of non-random mating were lower, with 81% of detection by *J*_RPTI_ against 74% by *J*_PTI_. However, in the case of the detection of sexual isolation, the *J*_RPSI_ test had 0% power while *J*_PSI_ had 74% (all the non-random mating detected cases).

As mentioned before, when we want to distinguish sexual isolation versus sexual selection effects, we need to identify the expectation *P*’_*Rij*_ of random mating from the frequencies in mated individuals. This expectation is formalized by equation (3) and *P*’_*Rij*_ is usually estimated by counting the different phenotypes within the matings.

However, a problem arises if we separate the counting in different mating rounds and there are very few mating per round. In such case, it is difficult to capture the marginal effect of each phenotype. Obviously, the individual encounter model is the extreme worst scenario since we only can assign a value of 1 for one of the matings at each round. Indeed, performing the estimation of *P*’_*Rij*_ in that way, under the individual encounter model, corresponded to the relative frequency of observed matings after the breeding season, i.e. *q*’_*Rij*_. Thus, the value of RPSI_*ij*_ was 1 and so *J*_RPSI_ was 0, which explains the observed 0% power for the *J*_RPSI_ test under the individual encounter model.

There are different ways to avoid this problem. Ideally, if we have independent estimates of the mutual propensities we can compute directly *P*’_*Rij*_ by (3). When we do so, the 100% of the effects were detected as sexual isolation by *J*_RPSI_ (i.e. 81% power).

Nevertheless we may not have the mutual propensity estimates and so, we should group the information of different rounds to obtain reliable estimates of *P*’_*Rij*_. As a rule of thumb, combining rounds so that the number of matings equals the product of the phenotypic classes (*k*_1_ × *k*_2_) should be enough to provide more power than the constant frequency test. Alternatively, we could use the constant frequency test that still provides a power of 74% for detecting sexual isolation even under the individual encounter model.

Therefore, the constant frequency test (*J*_PTI_) seems to be a good conservative control in this case, because it is not expected that the partition of *J*_PTI_ detect sexual isolation when the more powerful *J*_RPTI_ detects sexual selection instead. If that happens, we should concern about the *J*_RPSI_ estimate, especially if the number of matings per round is lower than *k*_1_ × *k*_2_.

### 3.3. Non-random mating: Sexual selection

Here we considered the same length polymorphism trait so that females with long length polymorphism had more mating vigor. The population size, mating rounds and initial frequencies were defined as before.

To model female sexual selection, the mutual propensities were different so that *L* females had three times more propensity than the *S* ones, so *m*_*LL*_ = *m*_*LS*_ = 3*m*_*SL*_ and *m*_*SS*_ = *m*_*SL*_. The specific value of the maximum propensity depended on the encounter model i.e. *m*_*LL*_ = 1 for the individual encounter and *m*_*LL*_ = 10 for the mass encounter model.

In Table 3 we see a specific mass encounter example, with significant *J*_RPTI_ (*J*_RPTI_ = 0.123, *p* - value = 0.007) but non-significant *J*_PTI_ (*J*_PTI_ = 0.006, *p* - value = 0.887). Sexual selection was also detected.

**TABLE 3.** Result of 15 mating rounds under mass encounter model with female sexual selection: *m*_*LL*_ = *m*_*LS*_ = 10 = 3*m*_*SL*_, *m*_*SS*_= *m*_*SL*_.

| Round $r$ | $p_{1rL}$ | $p_{2rL}$ | $L \times L$ | $L \times S$ | $S \times L$ | $S \times S$ |
| --- | --- | --- | --- | --- | --- | --- |
| 1 | 0.5 | 0.5 | 4 | 2 | 1 | 0 |
| 2 | 0.478 | 0.489 | 1 | 3 | 1 | 2 |
| 3 | 0.471 | 0.506 | 4 | 0 | 1 | 1 |
| 4 | 0.456 | 0.481 | 3 | 3 | 2 | 0 |
| 5 | 0.423 | 0.465 | 4 | 2 | 1 | 3 |
| 6 | 0.393 | 0.459 | 2 | 2 | 1 | 2 |
| 7 | 0.370 | 0.463 | 2 | 2 | 3 | 1 |
| 8 | 0.348 | 0.435 | 3 | 2 | 0 | 3 |
| 9 | 0.289 | 0.447 | 2 | 1 | 3 | 1 |
| 10 | 0.258 | 0.387 | 0 | 1 | 1 | 2 |
| 11 | 0.259 | 0.407 | 2 | 0 | 2 | 2 |
| 12 | 0.238 | 0.333 | 0 | 1 | 2 | 2 |
| 13 | 0.250 | 0.312 | 0 | 1 | 1 | 5 |
| 14 | 0.333 | 0.444 | 0 | 3 | 1 | 1 |
| 15 | 0.0 | 0.750 | 0 | 0 | 3 | 1 |
| Observed total |  |  | 27 | 23 | 23 | 27 |
| | | $q'_R$ | 0.27 | 0.23 | 0.23 | 0.27 |
| | | $q_R$ | 0.152 | 0.186 | 0.307 | 0.355 |
| | | $\Delta q_R$ | 0.118 | 0.044 | -0.077 | -0.085 |
| | | $Ln(q'_R/q_R)$ | 0.575 | 0.212 | -0.289 | -0.274 |
| $J_{RPTI}$ (p-value) | 0.123 (0.007) | $J_{RPSS}$ (p-value) | 0.075 (0.023) | | | |
| $J_{PTI}$ (p-value) | 0.006 (0.887) | | | | | |

$q'_R$ : observed mating frequencies; $q_R$ : expected random mating frequencies computed following the model in equation (1) by taking equal propensities; $\Delta q_R$ : $q'_R - q_R$ . $Ln$ : natural logarithm.

Noteworthy, the generalized indices had a lot of more power for the detection of the sexual selection non-random mating effects than the constant frequency indices. Under the mass-encounter model, the percentage of non-random mating (*J*_*RPTI*_) was 73%, with 59% of female sexual selection, 3% male sexual selection, and 5% of sexual isolation. The constant frequency methods however were not able of detecting non-random mating effects (1% of detection by *J*_PTI_).

The results under the individual model were similar, the constant indices had no power (0.8%) and the new ones detected non-random mating with female sexual selection 60% of the time.

## 4. Discussion

In a previous work we had described the change in the mating phenotypes due to mate choice as the gain of information with respect to random mating. That model is valid for polygamous species, or even for monogamous when only a fraction of the population is involved in the breeding season. In such cases, the processes of the encounter and mating correspond to sampling with replacement of the mating phenotypes from the population (Carvajal-Rodríguez, 2018).

In the current work we have generalized the previous model to incorporate the case of monogamous species with small population size, so that the mating process resembles a sampling without replacement.

Notably, the change in the phenotypes can still be described by the flow of information between the random and non-random mating cases and it can be partitioned in the sum of the information due to sexual isolation and sexual selection, plus a mixed effect term. The new formulas become the same as for the corresponding constant ones as soon as the population frequency values are set to constant.

Before obtaining the new information indices we previously defined a generalized version of the PTI, PSI and PSS pairwise statistics (Rolán-Alvarez and Caballero, 2000). The new statistics measure the same as the previous ones (total change, sexual isolation and sexual selection, respectively) without requiring that the population frequencies are constant.

When the frequencies vary over different mating rounds, the tests performed by the new indices have more power than the constant ones which are more conservative. The difference becomes extreme for the sexual selection scenarios. In this case, the indices that assume constant frequency have almost no detection power, while the generalized indices have up to 70%. The reason is that the variation of population frequencies over the mating rounds, hides the sexual selection effect at the end of the breeding season. This happens because in the first rounds most of mating favour some phenotypes (e.g. females *L*) so that the encounters and mating for such phenotypes diminish at the end of the season.

We have tested the extreme case of monogamous species with low population size where all adults perform the mating. As the proportion of mating individuals decreases, the differences between the two kinds of indices disappear.

Last but not least, the application of the new statistics requires the estimation of frequencies at each mating round, which can be difficult especially under individual encounter-mating models. Simulations show that in the sexual isolation scenario, the previous indices still perform well under variable population frequencies and so, the use of one or another type of statistics will depend on the biological scenario as well as in the availability and easiness to split the sampling in more than one mating round.

## Acknowledgement

I would like to thank Carlos Canchaya for their valuable comments on the manuscript. This work was supported by Xunta de Galicia (Grupo de Referencia Competitiva, ED431C2016 - 037), Ministerio de Economía y Competitividad (CGL2016 - 75482 - P) and by Fondos FEDER (“Unha maneira de facer Europa”).

